# Electrophysiological validation of premotor interneurons monosynaptically connected to the aCC motoneuron in the *Drosophila* larval CNS

**DOI:** 10.1101/2020.06.17.156430

**Authors:** Carlo N. G. Giachello, Aref Arzan Zarin, Hiroshi Kohsaka, Yuen Ngan Fan, Akinao Nose, Matthias Landgraf, Richard A. Baines

**Affiliations:** Division of Neuroscience and Experimental Psychology, School of Biological Sciences, Faculty of Biology, Medicine and Health, University of Manchester, Manchester Academic Health Science Centre, Manchester, M13 9PL, UK; Department of Biology, Texas A&M, University, College Station, United States; Department of Complexity Science and Engineering, Graduate School of Frontier Sciences, University of Tokyo, Tokyo, Japan; Department of Zoology, University of Cambridge, Cambridge, CB2 3EJ, UK

## Abstract

Mapping the wired connectivity of a nervous system is a prerequisite for full understanding of function. In this respect, such endeavours can be likened to genome sequencing projects. These projects similarly produce impressive amounts of data which, whilst a technical *tour-de-force*, remain under-utilised without validation. Validation of neuron synaptic connectivity requires electrophysiology which has the necessary temporal and spatial resolution to map synaptic connectivity. However, this technique is not common and requires extensive equipment and training to master, particularly when applied to the small CNS of the *Drosophila* larva. Thus, validation of connectivity in this CNS has been more reliant on behavioural analyses and, in particular, activity imaging using the calcium-sensor GCaMP. Whilst both techniques are powerful, they each have significant limitations for this purpose. Here we use electrophysiology to validate an array of driver lines reported to label specific premotor interneurons that the *Drosophila* connectome project suggests are monosynaptically connected to an identified motoneuron termed the anterior corner cell (aCC). Our results validate this proposition for four selected lines. Thus, in addition to validating the connectome with respect to these four premotor interneurons, our study highlights the need to functionally validate driver lines prior to use.

## Introduction

A long appreciated advantage of using invertebrates for neuroscience research is the existence of ‘identified neurons’ which, as their name implies, can be uniquely identified and returned to across preparations. The ability to record from neurons in *Drosophila* has made particular use of this and has combined cell-specific electrophysiology with genetics (Baines and Bate, 1998; Baines et al., 1999; Baines et al., 2001; Baines et al., 2002; Worrell and Levine, 2008; Ryglewski et al., 2012; Srinivasan et al., 2012a, b; Kadas et al., 2017). More recently, the first instar larval *Drosophila* nervous system has been reconstructed using serial section transmission electron microscopy (ssTEM) (Schneider-Mizell et al., 2016; Gerhard et al., 2017; Larderet et al., 2017; Saumweber et al., 2018; Zarin et al., 2019) and annotated into a CATMAID dataset (Saalfeld et al., 2009). The eventual goal of this endeavour is to identify and characterise all neurons in the larval CNS and, moreover, to trace their wired synaptic connectivity. Once identified, specific neurons are amenable to genetic manipulation using a range of driver lines, including specific split-GAL4 lines (Kohsaka et al., 2014; Fushiki et al., 2016; Hasegawa et al., 2016; Schneider-Mizell et al., 2016; Kohsaka et al., 2019). Thus, a large dataset of putative neuronal connections is now available and researchers are making increasing use of this information and derived driver lines to investigate the physiology of the larval CNS.

However, there is a danger of ‘running before walking’ when utilizing this impressive dataset. This is because connectivity between neurons has rarely been directly verified, but inferred from EM and validated through behavioural analysis and/or functional imaging using GCaMP to establish correlated activity (Kohsaka et al., 2019; Zarin et al., 2019). A caveat when using behavioural responses, and to a lesser extent functional imaging, is that these methods can lack the required resolution necessary to establish unambiguously whether or not two cells are monosynaptically connected; as opposed to being connected through intermediate neurons (i.e. polysynaptic). A combination of optogenetics and Ca^2+^-imaging, together with pharmacology to block action potential firing, has been used to infer monosynaptic connectivity (Sales et al., 2019), but again analysis can be complicated by variability in response recorded in the postsynaptic neuron. By contrast, electrophysiology provides what has often been described as the ‘gold-standard’ to verify functional connectivity between neuron pairs. Whilst electrophysiology has been used to validate the connectome (Zwart et al., 2016), its use remains very limited. This is undoubtedly due to the technical nature of electrophysiology and the paucity of researchers that are able to employ this approach.

We have, in this study, adopted a traditional electrophysiological approach to screen a range of interneuron driver lines that have been suggested to monosynaptically-connect with a specific motoneuron, termed the anterior corner cell (aCC). Because of its larger soma and midline-dorsal location, this cell has been the subject of numerous studies and remains a key focus of many groups (Worrell and Levine, 2008; Ryglewski et al., 2012; Srinivasan et al., 2012a, b; Giachello and Baines, 2015; Giachello et al., 2016). We confirm that aCC is monosynaptically driven by two excitatory premotor cholinergic interneurons (A27h and A18a/CLI2), in addition to two GABAergic premotor interneurons, A23a and A31k. Whilst this validates the connectome, we observe that multiple driver lines, indicated to drive gene expression in A23a or A31k, show dramatically different effects, with some not effective at all. This underlies the concern of using driver lines without a reliable method to establish functional connectivity to the neuron being investigated.

## Materials and Methods

### Drosophila rearing and stocks

All *Drosophila melanogaster* strains were grown and maintained on standard corn meal medium at 25°C. To optogenetically manipulate neurons we used the following transgenic lines: *ChR;NaChBac* (*w*; 20xUAS-T159C-ChR2; UAS-NaChBac-EGFP / TM6C*^*Sb,Tb*^) which was created by crossing *y*^*1*^,*w*; 20xUAS-T159C-ChR2; Dr*^*1*^*/ TM6C*^*Sb,Tb*^ (#52258, Bloomington Drosophila Stock Center, Indiana, USA) and *UAS-NaChBac-EGFP/TM3*^*Sb*^ (#9467, BDSC). Chrimson: *w*^*1118*^; *20xUAS-IVS-CsChrimson*.*mVenus;* + (#55135, BDSC) and *w*; UAS-H134R-ChR2;* + (a gift from Stefan Pulver). Interneuron expression driver lines used were: *w*^*1118*^; +; *R36G02-Gal4* (A27h-Gal4, #49939, BDSC) expresses in the premotor interneuron A27h, in addition to a few other neurons (Fushiki et al., 2016), *w*^*1118*^; +; *47E12-Gal4* (*CLIs-Gal4*, #50317, BDSC) which expresses in CLI1 and CLI2 and some unknown interneurons (Hasegawa et al., 2016). *w*^*-*^; +; *R47E12-Gal4; cha3*.*3-Gal80* (CLI1/2-Gal4) which expresses in CLI1 and CLI2 (Hasegawa et al., 2016). *w*^*-*^; *tsh-Gal80; R47E12-Gal4; cha3*.*3-Gal80* (CLI1-Gal4,) which is specific for CLI1 (Hasegawa et al., 2016). *w*^*-*^; +; *R15B07-Gal4* (CLI2-Gal4 or A18a-Gal4) which is specific for CLI2. *w*^*-*^; +; *Gad1-T2A-Gal4 / TM6b* which expresses in all GABAergic neurons. Three different lines designed to target A23a: *R78F07-Gal4* (Zarin et al 2019), *R78F07-AD; R49C08-DBD* split Gal4 and *R41G07-AD; R78F07-DBD* split Gal4 (also termed *SS04495-Gal4*) (Kohsaka et al., 2019). Three different lines designed to target A31k: *R87H09-Gal4* (Zarin et al 2019), *R20A03-AD; R87H09-DBD* split Gal4 and *R20A03-AD* (Zarin, unpublished); *R93B07-DBD* split Gal4 (also termed *SS04399-Gal4*) (Kohsaka et al., 2019).

### Electrophysiology

Electrophysiological recordings were performed in third (L3) and first (L1) instar larvae as previously described (Baines et al., 1990; Marley and Baines, 2011). aCC motoneurons were identified in bright-field microscopy, while GFP-expression driven by A27h-Gal4 was used to recognise A27h interneurons. Whole-cell voltage- and current-clamp recordings were achieved using thick-walled borosilicate glass electrodes (GC100F-10, Harvard Apparatus, Edenbridge, UK) fire polished to resistances of 10-15 MΩ (L3 aCC) and 15-20 MΩ (L1 aCC and L3 A27h). Recordings were made using a Multiclamp 700B amplifier controlled by pCLAMP (version 10.4) via a Digidata 1440A analog-to-digital converter (Molecular Devices, Sunnyvale, CA). Traces were sampled at 20 kHz and filtered online at 10 kHz. Cells with input resistance <0.5 GΩ were not considered for analysis. External saline composition was as follows: 135 mM NaCl, 5 mM KCl, 4 mM MgCl_2_·6H_2_O, 2 mM CaCl_2_·2H_2_O, 5 mM TES and 36 mM sucrose, pH 7.15. Internal patch solution was as follows: 140 mM K^+^-D-gluconate, 2 mM MgCl_2_·6H_2_O, 2 mM EGTA, 5 mM KCl, and 20 mM HEPES, pH 7.4. KCl, CaCl_2_, MgCl_2_ and sucrose were purchased from Fisher Scientific (Loughborough, UK); all remaining chemicals were obtained from Sigma-Aldrich (Poole, UK). Optogenetic stimulation of ChR was achieved by using a λ470 LED (bandwidth 25 nm, irradiance 15.62 mW·cm^-2^; OptoLED, Cairn Instruments, Kent, UK) connected to an Olympus BX51WI microscope (Olympus Corporation, Tokyo, Japan). Light was controlled by Clampex (version 10.4) and pulsed for 1 second during recordings. The same stimulation protocol was also applied to Chrimson and H134R-ChR, in Supplementary Figure 1. Chrimson is designed to be a red-shifted version of channelrhodopsin, but still strongly responds to blue light (at least in our isolated CNS preparation). Therefore, we decided to use the same light source in all our experiments to minimise variations in light irradiance. In order to assess NaChBac functionality, the construct ChR; NaChBac was expressed under the control of A27h-Gal4. A27h interneurons were recorded in current-clamp mode, holding their membrane potential at -60 mV and optogenetically stimulated as described above. Cells were recorded in both absence and presence of TTX (2 μM; Alomone Labs, Israel) to block endogenous sodium voltage-dependent channels and isolate the NaChBac conductance. NaChBac activation was also examined by injecting constant current into A27h in presence of 2 μM TTX. In this experiment, A27h was held at -90 mV and incremental pulses of 1 pA, from -5 to +45pA / 500 ms-long, were applied. Threshold was defined as the voltage at the onset of each spike and assigned by careful visual inspection of the raw data. NaChBac steady-state inactivation was analysed by repeating the optogenetic stimulation protocol (λ470 nm, 1 s), holding A27h at different potentials (−90, -60, -40 and -20 mV). Since NaChBac kinetics are faster than 1 s (light pulse), the contribution of ChR activation was excluded by measuring NaChBac amplitude from the maximum value of the peak until the signal fully decays once the peak has occurred. Amplitudes were then plotted against the A27h holding membrane potential.

**Figure 1.**
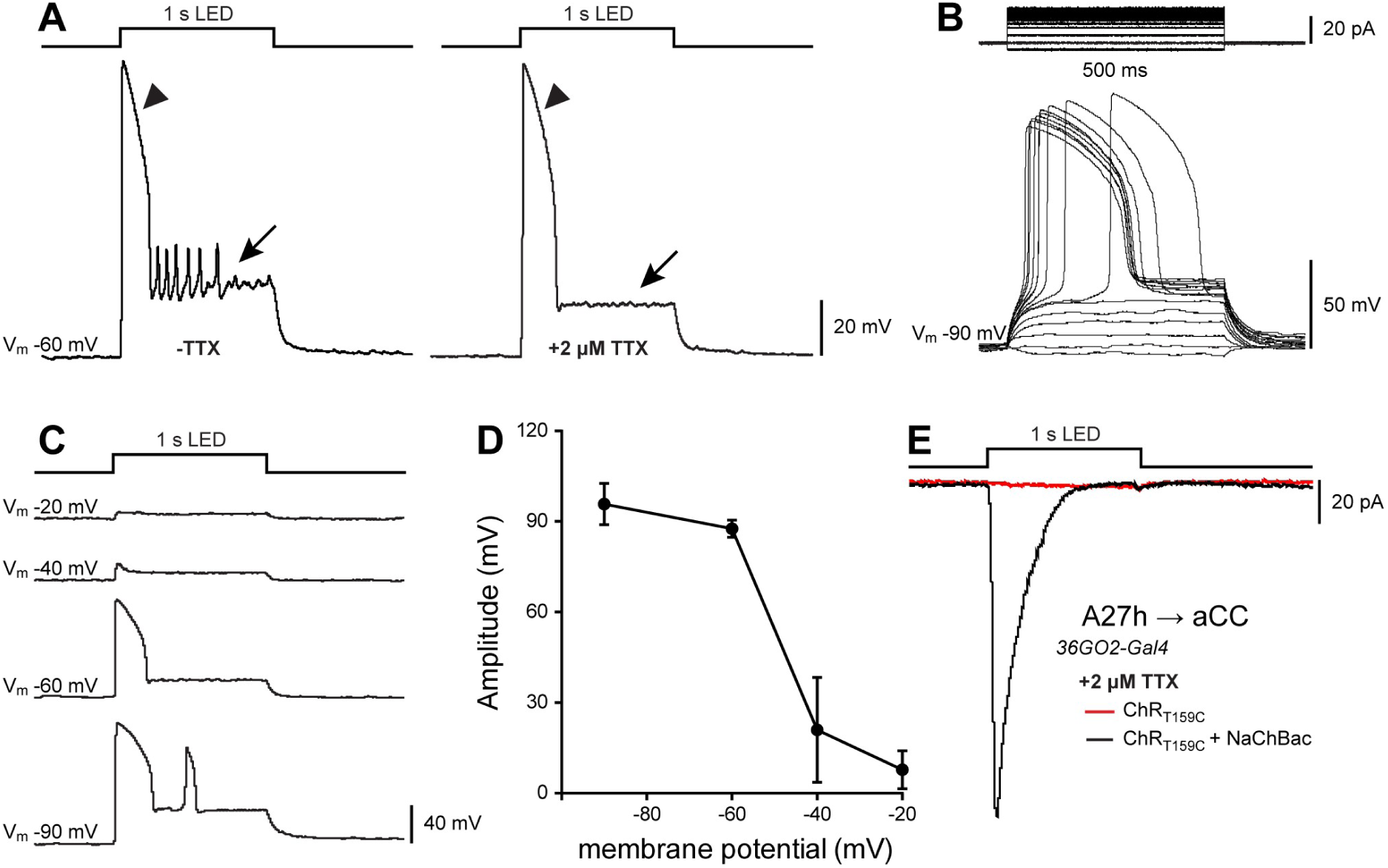
*ChR; NaChBac* is a powerful tool to verify monosynaptic connectivity. (A) A representative current-clamp recording from the A27h interneuron overexpressing both ChR and NaChBac. Optogenetic stimulation (λ470 nm, 1 s) induced activation of NaChBac which persists in presence of 2 µM TTX (arrowheads). Conversely, APs produced by activation of endogenous voltage-gated sodium channels were blocked after TTX application (arrows). (B) Voltage dependence of NaChBac activation recorded from A27h in current-clamp. A27h depolarisation was elicited by injecting constant current steps (1 pA steps/0.5 s) in the presence of TTX. (C-D) Voltage-dependent inactivation of NaChBac. Peak amplitude was recorded and measured from A27h held at different voltages (from -90 to -20 mV) during optogenetic stimulation (λ470 nm, 1 s). NaChBac activation is affected at potentials more positive than -40 mV. Note: there is a second activation (peak) of NaChBaC at -90mV. (D) Averaged data ± SEM (*n* = 3). (E) Sample recording of synaptic drive to aCC, recorded in voltage-clamp, following optogenetic activation of A27h (λ470 nm, 1 s). In presence of TTX, co-expression and activation of both ChR and NaChBac in A27h produced a clear synaptic input in aCC (inward current, black trace), thus confirming the existence of a monosynaptic connection between these two neurons. As a control, TTX successfully blocked aCC inputs when only ChR, but not NaChBac, was expressed in A27h (red trace).

In order to assess monosynaptic connectivity between aCC and putative presynaptic interneurons, the construct ChR; NaChBac was expressed under the control of interneuron type-specific Gal4 expression lines. Some of these lines do not express transgenes homogenously in every segment. To ensure that our recordings were performed from the correct hemisegment, we exploited the EGFP-tag on NaChBac, quickly verifying its expression before each data acquisition. aCC motoneurons were recorded in whole-cell patch clamp in absence and presence of 2 μM TTX. Optogenetic stimulation (λ470 nm, 1 s) was repeated 5 times per cell. Traces were averaged and examined using Clampfit (version 10.4). Synaptic connections were examined in both current and voltage clamp, by holding aCC at -60mV to visualise excitatory inputs (A27h, CLIs). Inhibitory connections (all GABAergic, A23a, A31k) were also visualised by holding aCC at -40 mV, further away from the chloride reversal potential. To measure the amplitude of inputs, the change from baseline to peak amplitude was determined. Currents shown were normalized for cell capacitance (determined by integrating the area under the capacity transient resulting from a step protocol from -60 to -90 mV). Inhibition was also quantified as a decrease in action potential firing by the postsynaptic cell. Action potentials were evoked by injecting a supra-threshold continuous depolarising current into aCC. Optogenetic stimulation (λ470 nm, 1 s) was repeated 5 times per recording. Action potentials were counted over a time window of 1 s before, during and after optogenetic stimulation, once firing was resumed.

### Drugs

Synaptic transmission was disrupted by blocking endogenous sodium channels with 2 μM TTX (Alomone Labs, Israel). Inhibitory inputs were selectively blocked with either 10 µM PTX (Sigma, UK) or 1 mM Gabazine (SR95531, Sigma, UK), an antagonist of the *Drosophila* GABA_A_ receptor Rdl. Cholinergic inputs were selectively blocked with 1 mM mecamylamine (Sigma, UK). All drugs were bath applied during electrophysiological recording.

### Statistical Analysis

Data was acquired and imported into Microsoft Excel (Microsoft Corp., Redmond, WA). All data are presented as the mean ± SEM. No statistical test was used on the measurement of synaptic inputs because of the descriptive nature of these data (no comparisons among groups). Conversely, statistical analyses were conducted on firing plots, to measure the premotor inhibitory drive, by using repeated measures one-way ANOVA followed by Bonferroni’s *post-hoc* multiple comparisons test. The null hypothesis was rejected at the 0.05 level. *P* values < 0.05 were considered significant. Significance was shown as * = *P* < 0.05, ** = *P* < 0.01, and not significant values were not noted. Statistical tests were performed in GraphPad Prism (version 7, GraphPad Software, San Diego, CA). Figures were assembled with Adobe Illustrator CS3 (Adobe, San Jose, CA, USA).

## Results

### A27h is monosynaptically connected to aCC

To verify monosynaptic connectivity between two neurons, we adopted a recently described method - Tetrodotoxin (TTX) Engineered Resistance for Probing Synapses (TERPS) (Zhang and Gaudry, 2018) - which exploits the insensitivity of the voltage-gated sodium channel (Na_v_) from *Bacillus halodurans*, called NaChBac, to tetrodotoxin (TTX) (Ren et al., 2001). We generated a transgenic stock containing both the T159C variant of channelrhodopsin (T159C-ChR) (Berndt et al., 2011) and NaChBac, to allow both of these Gal4 responsive UAS-transgenes to be expressed simultaneously. Expressing both of these transgenes in a presynaptic interneuron of choice allows monosynaptic connectivity to be established by optogenetically activating the interneuron whilst patch recording from a presumed postsynaptic cell. Persistence of synaptic drive in the presence of TTX, which blocks spiking activity in all neurons with the exception of those expressing the TTX-insensitive bacterial NaChBac, proves monosynaptic connectivity.

We piloted this approach using the A27h neuron, a well-characterised cholinergic premotor interneuron that the connectome indicates is synaptically connected to aCC. This connectivity has been confirmed by paired whole-cell recordings (Fushiki et al., 2016). We performed electrophysiological recordings from A27h interneurons where both ChR and NaChBac overexpression was driven by *R36G02-Gal4*, also termed ‘*A27h-Gal4* ‘(Fushiki et al., 2016). As expected, depolarisation of A27h via optogenetic stimulation (ChR, λ470 nm, 1 s) induced a large and slow-inactivating depolarisation, due to the activation of NaChBac, which persisted in presence of 2 µM TTX (arrowheads in Figure 1A). However, endogenous action potential firing in A27h (arrows in Figure 1A) is, as expected, absent in the presence of TTX (Figure 1B). Depolarisation of A27h via injection of constant current (1 pA steps/0.5 s) directly into A27h also produced NaChBac activation which occurred with an activation threshold of -65 ± 12 mV (Figure 1B) in the presence of TTX. The rate of inactivation of NaChBac increases steeply as a function of voltage (Ren et al., 2001). We therefore determined NaChBac steady-state inactivation by measuring the peak amplitude, activated by ChR (λ470 nm, 1 s), following changes to A27h membrane potential (Figure 1C). We observed that at membrane potentials more positive than -40 mV, NaChBac activation is severely reduced (Figure 1D), similar to the detailed descriptions of NaChBac properties when expressed in CHO-K1 cells (Ren et al., 2001). This suggests that NaChBac inactivation, more severe at relatively depolarised membrane potentials, could potentially be a limitation of this method, and this should be taken into account when planning and interpreting experiments (see Discussion).

Finally, once we had established that NaChBac is active in A27h, we confirmed that A27h and aCC are monosynaptically connected, by optogenetic stimulation of A27h and simultaneously recording from aCC motoneurons. In the presence of TTX (2 μM), we observed a clear response in aCC (Figure 1E, *R36G032-Gal4>T159C-ChR; NaChBac*: -20.78 ± 4.01 pA/pF, *n* = 5). Conversely, TTX completely blocked aCC synaptic drive in the absence of NaChBac (−0.07 ± 0.01 pA/pF, *n* = 5). These results corroborate the use of ChR combined with NaChBac to identify monosynaptically-connected neuron pairs. In the next set of experiments, we use this tool to validate the identification of additional aCC synaptic partners, identified from the connectome, but not yet verified by electrophysiology.

### A18a is monosynaptically connected to aCC

We tested two additional cholinergic pre-motor interneurons termed A18a and A18b3. The connectome shows that A18a, but not A18b3, is monosynaptically connected to aCC (Zarin et al., 2019). These cells are cholinergic lateral interneurons (originally termed CLIs 2 and 1, respectively) (Hasegawa et al., 2016). Four different Gal4 expression lines were tested: *R47E12-Gal4*, also termed ‘*CLIs-Gal4’*, targets several unknown interneurons including A18a and A18b3 and potentially also sensory neurons; *R47E12-Gal4; cha3*.*3-Gal80*, also termed ‘*CLI1/2-Gal4’*, restricts expression to A18a and A18b3 (Hasegawa et al., 2016); *tsh-Gal80; R47E12-Gal4, cha3*.*3-Gal80*, also called ‘*CLI1-Gal4’*, is selective for CLI1/A18b3 only (Hasegawa et al., 2016); and *R15B07-Gal4*, also termed ‘*CLI2-Gal4’*, is specific for CLI2/A18a (Zarin, unpublished).

Figure 2 shows ChR-evoked inputs to aCC recorded in absence (Figure 2A) and presence (Figure 2B) of TTX. The *CLI-Gal4* line, less specific compared to the other ones, exhibited the largest synaptic drive to aCC, which is notably reduced, but not fully blocked, by the presence of TTX (−9.55 ± 3.19 pA/pF, *n* = 4, to -3.57 ± 0.92 pA/pF, *n* = 6, Figure 2A-C).

**Figure 2.**
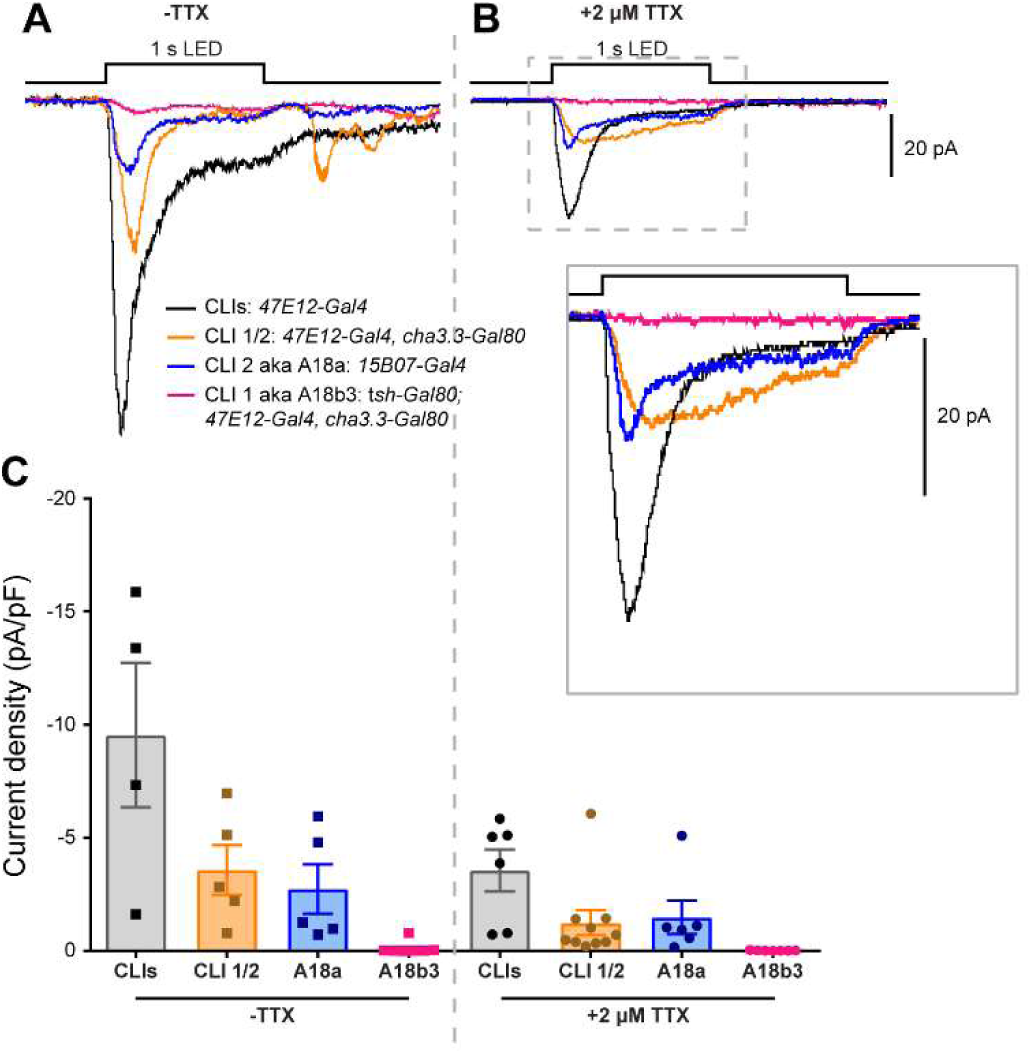
A18a/CLI2 and aCC are monosynaptically connected. (A-B) Excitatory synaptic inputs to aCC recorded in absence (A) and presence of TTX (B and inset). Four different Gal4 lines were tested to drive ChR and NaChBac expression in the cholinergic lateral interneurons 1 and 2 (CLI 1 and 2). (C) Quantification of aCC synaptic drive revealed that CLI2, but not CLI 1, is monosynaptically connected to aCC. The TTX-induced reduction of synaptic current amplitudes (e.g. *CLIs-Gal4*) suggests the existence of additional unknown intermediary neurons located upstream of aCC and activated by neurons expressing ChR; NaChBac.

This reduction is probably ascribed to the TTX-dependent block of unknown intermediary neurons upstream of aCC which, in turn, were activated by neurons co-expressing ChR and NaChBac. A similar decrease, although less pronounced, was observed with *CLI1/2-Gal4* line (from -3.59 ± 1.10 pA/pF, *n* = 5, to -1.25 ± 0.55 pA/pF, *n* = 10) and *CLI2-Gal4* (from -2.74 ± 1.09 pA/pF, *n* = 5, to -1.49 ± 0.73 pA/pF, *n* = 6). Interestingly, in the presence of TTX, both *CLI1/2-Gal4* and *CLI2-Gal4* lines showed comparable amplitudes, suggesting a major contribution from CLI2 (A18a) compared to CLI1 (A18b3). Therefore, we also tested *CLI1-Gal4* line and observed no detectable synaptic drive to aCC, either in the absence or presence of TTX (−0.09 ± 0.09 pA/pF, *n* = 8, to -0.01 ± 0.01 pA/pF, *n* = 9). Thus we conclude that CLI2 (A18a), but not CLI1 (A18b3), synaptically drives aCC and, moreover, does so monosynaptically. This is in full agreement with the connectome (Zarin et al., 2019).

### aCC receives GABAergic inputs

The connectome identifies inhibitory neurons making direct synaptic connections to motoneurons, including aCC (Clark et al., 2018, Kohsaka, 2019 #2020). We found that expression of NaChBac in all GABAergic neurons (*Gad1-T2A-Gal4*), as well as all cholinergic neurons (*ChAT-BAC-Gal4*), is lethal. Thus, we initially expressed just T159C-ChR in GABAergic neurons in order to visualise the total inhibitory synaptic input to aCC. We recorded aCC motoneurons in current-clamp, injecting a supra-threshold depolarising current to elicit action potentials (APs, motoneurons are not spontaneously active). Activation of GABAergic neurons (λ470 nm, 1s) produced a significant decrease in AP firing in aCC for the duration of the light pulse (8.89 ± 1.85 to 0.35 ± 0.24, *n* = 7, *P* = 0.0103, repeated measures one-way ANOVA followed by Bonferroni’s post hoc test, Figure 3A). Firing quickly resumed following cessation of light stimulation (4.26 ± 2.75 *vs*. 8.89 ± 1.85, *n* = 7, *P* = 0.1166). As further evidence for inhibitory drive, voltage-clamp traces showed increasing synaptic current density as aCC was depolarised away from the chloride reversal potential. A current of +0.25 ± 0.13 pA/pF (*n* = 8) at -60 mV increased, as expected for Cl^-^ ions, to +1.65 ± 0.36 pA/pF (*n* = 8) in the same cells at -40 mV (Figure 3B-C). Similarly, current-clamp recordings clearly exhibited an increasing hyperpolarising drive to aCC of -1.93 ± 0.80 mV and -6.56 ± 1.18 mV, at -60 mV, or -40 mV, respectively (Figure 3D-E).

**Figure 3.**
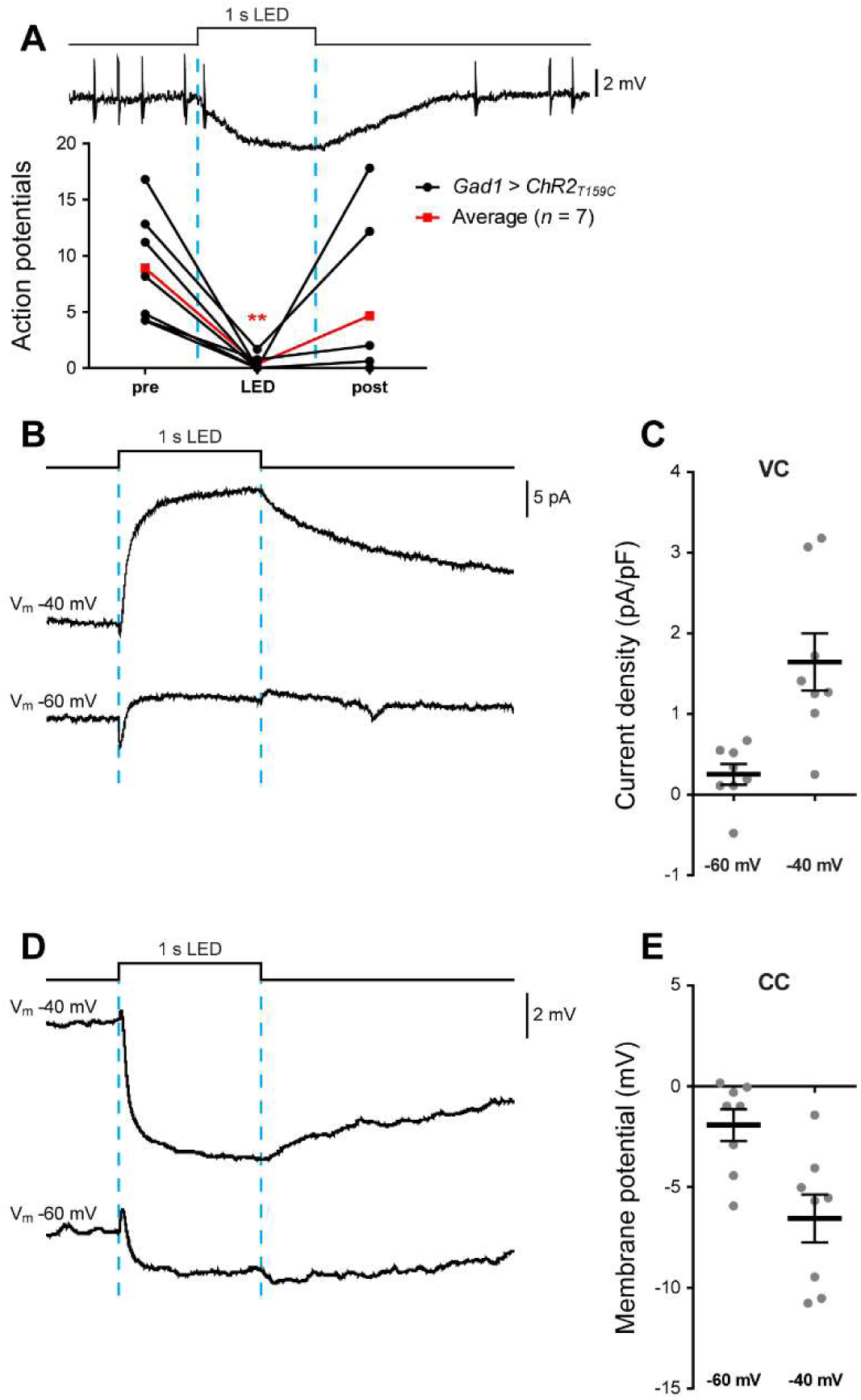
aCC motoneurons receive inputs from GABAergic interneurons. (A) Sample trace and quantification of APs evoked by injecting a supra-threshold depolarising current into aCC. Optogenetic activation of all GABAergic interneurons (*Gad1-T2A-Gal4>UAS-T159C-ChR*, λ470 nm, 1 s) almost completely inhibited AP firing in aCC (F_(2, 18)_ = 8.391, *P* = 0.0137, repeated measures one-way ANOVA, *n* = 7, black lines), clearly showing that aCC receives inhibitory inputs. Average values are shown in red. (B-C) Sample trace and quantification of the inhibitory drive to aCC recorded in voltage-clamp mode. The same cells were recorded at holding potentials of -40 and -60 mV. As expected, we observed a large outward current (at -40 mV) which attenuated at more negative potentials (−60 mV) close to the chloride reversal potential (approx. -70 mV). (D-E) Sample trace and quantification of the inhibitory drive to aCC recorded in current-clamp mode showing a clear hyperpolarisation of aCC and same attenuation at -60 mV compared to -40 mV.

### A23a is monosynaptically connected to aCC

The connectome identifies A23a as a GABAergic interneuron directly presynaptic to aCC, involved in locomotion and activated in both forward and backward peristaltic waves (Kohsaka et al., 2019). We tested three different lines reported to drive Gal4 in the A23a interneuron: *R78F07-Gal4* (Zarin et al., 2019); *R78F07-AD; R49C08-DBD* split Gal4 (Zarin, unpublished); and *R41G07-AD 78F07-DBD* split Gal4, also termed *SS04495-Gal4* (Kohsaka et al., 2019).

Optogenetic activation of *R78F07-Gal4* expressing neurons produced a decrease in AP firing in aCC consistent with an inhibitory input (4.70 ± 1.07 *vs*. 1.32 ± 0.61 APs, *n* = 6, *P* = 0.0406, repeated measures one-way ANOVA followed by Bonferroni’s post hoc test; post LED = 5.02 ± 1.01 APs, Figure 4A). However, *R78F07-Gal4* exhibits an expression pattern that is more diverse than expected: targeting Gal4 to additional cells located on the ventral surface of the ventral nerve cord. This lack of specificity is reflected in the heterogeneity of responses recorded from aCC. In the absence of TTX, inhibitory inputs prevailed (input average: -1.73 ± 1.02 mV at -40 mV, *n* = 7, Figure 4B), whilst recordings performed in the presence of TTX exhibited an additional excitatory component (5 out of 8 cells), suggesting that the inhibitory input is polysynaptic and, further, that this driver also expresses in monosynaptically-connected excitatory premotor interneurons (input average: -0.46 ± 0.75 mV at - 40 mV, *n* = 10, Figure 4B). For example, Figure 4C shows evidence for a biphasic connection, where the inhibitory component seems to reliably occur in the absence of TTX, but which reverts to excitation after TTX application. These data suggest that *78F07-Gal4* is not selective, nor reliable, for activation of A23a.

**Figure 4.**
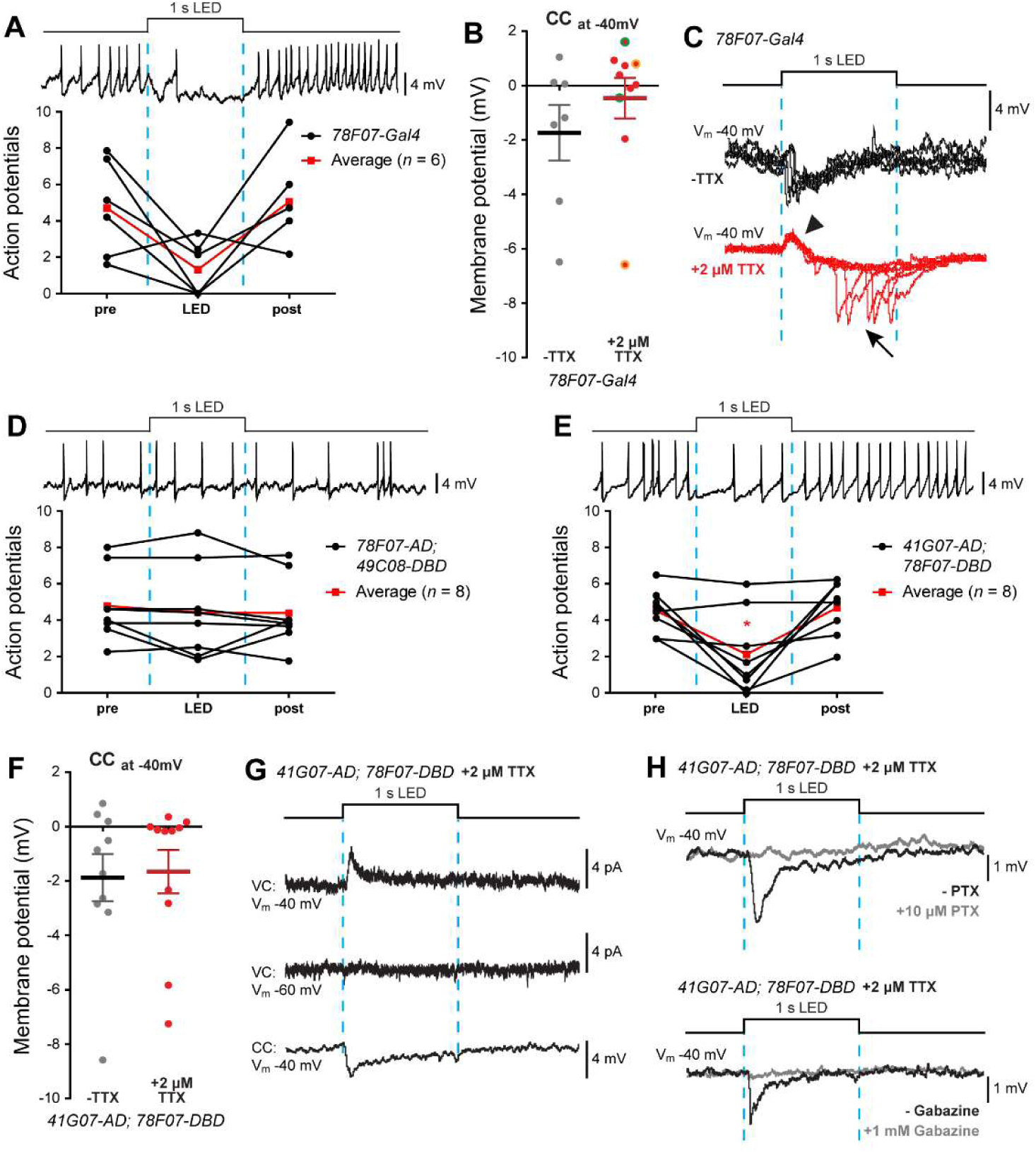
A23a and aCC are monosynaptically connected. (A) Optogenetic activation of *R78F07-Gal4* driving ChR; NaChBac reduces action potential firing in aCC (elicited by injection of constant current). On average, we observed an inhibitory effect (F_(2, 15)_ = 5.005, *P* = 0.0216, repeated measures one-way ANOVA, *n* = 6, black lines). Average values are shown in red. (B) Quantification of the synaptic inputs recorded from aCC (held at -40 mV) following optogenetic activation of *R78F07-Gal4* driving ChR; NaChBac, before and after 2 µM TTX application. In the presence of TTX, we observed a heterogeneous range of inputs with excitation prevailing over inhibition, thus suggesting a poor specificity for this line to target the GABAergic A23a interneuron. Some recordings (2 out of 8 cells) showed a biphasic connection where both the excitatory and inhibitory components were observed in the same cell (values highlighted with a different colour, +TTX group). (C) Raw electrophysiological sweeps from an example of a biphasic connection obtained with optogenetic activation of *R78F07-Gal4*. The same cell was recorded 5 times during optogenetic stimulation before (black traces) and after (red traces) TTX exposure. Whilst the inhibitory component seems to prevail before applying TTX, isolation of NaChBac -overexpressing neurons resulted in a reliable excitatory component (arrowhead) followed by a delayed erratic inhibitory component (arrow). (D) Optogenetic activation of *R78F07-AD; R49C08-DBD* split Gal4 did not affect aCC firing (F_(2, 21)_ = 0.8322, *P* = 0.4005, repeated measures one-way ANOVA, *n* = 8, black lines). Average values are shown in red. (E) Optogenetic activation of *R41G07-AD; R78F07-DBD* split Gal4 significantly reduced aCC firing (F_(2, 21)_ = 9.662, *P* = 0.0141, repeated measures one-way ANOVA, *n* = 8, black lines). Average values are shown in red. (F) Quantification of the synaptic drive to aCC (held at -40 mV) following optogenetic activation of *R41G07-AD; R78F07-DBD* split Gal4 in the absence, or presence, of 2 µM TTX. The prevalence of inhibitory inputs suggests a better specificity for this line in targeting A23a compared to the previous tested lines. (G) Sample traces showing the optogenetic activation of *R41G07-AD; R78F07-DBD* split Gal4, driving ChR; NaChBac. aCC were recorded both in voltage-(both at -60 and -40 mV) and in current clamp (at -40 mV) in presence of TTX. (H-I) Sample traces confirming that the A23a→ aCC synapse is GABAergic. Cells were recorded, as previously described, before (black trace) and after (gray trace) bath application of 10 µM Picrotoxin (H) or 1 mM Gabazine (I), two blockers of the *Drosophila* GABA_A_ receptor. In both cases, aCC inputs were abolished.

Next, we tested two different split Gal4 lines reportedly more specific in targeting Gal4 activity to the A23a interneuron. We started with *R78F07-AD; R49C08-DBD*, which has a very sporadic expression pattern, targeting A23a neurons in only a few hemisegments in any one animal. Optogenetic activation of *R78F07-AD; R49C08-DBD* did not significantly affect evoked AP firing in aCC (4.78 ± 0.69 *vs*. 4.42 ± 0.90 APs, *n* = 8, *P* > 0.99, repeated measures one-way ANOVA followed by Bonferroni’s post hoc test; post LED = 4.39 ± 0.68 APs, Figure 4D). Moreover, optogenetic activation utilising this driver did not exhibit a detectable input to aCC, tested under voltage-clamp at both -60 mV and -40 mV (data not shown). To exclude the possibility that our optogenetic activation is not sufficient to activate A23a, we additionally used Chrimson, a much more powerful ChR variant largely employed in behavioural studies (Klapoetke et al., 2014). Surprisingly, Chrimson-mediated activation exhibited a strong cholinergic input to aCC (Figure S1A), clearly confirming that *R78F07-AD; R49C08-DBD* does not target Gal4 activity selectively to GABAergic A23a.

For the other split-Gal4 line *SS04495-Gal4*: *R41G07-AD; R78F07-DBD* (Kohsaka et al., 2019), optogenetic activation produced an expected and significant reduction in evoked AP firing in aCC (from 4.50 ± 0.42 to 2.16 ± 0.80 APs, *n* = 8, *P* > 0.0415, repeated measures one-way ANOVA followed by Bonferroni’s post hoc test; post LED = 4.70 ± 0.54 APs, Figure 4E). Using this split-Gal4 line to drive *ChR; NaChBac* produced a reliable hyperpolarising drive to aCC that was unaffected by application of TTX (−1.88 ± 0.87 mV, *n* = 10, *vs*. -1.66 ± 0.80 mV at -40mV, *n* = 11, -TTX *vs*. +TTX, *P* = 0.8502, *t*-test, Figure 4F), showing inhibitory monosynaptic connectivity. To validate that the hyperpolarisation was due to the movement of Cl^-^ ions, we manipulated the membrane potential of aCC. Figure 4G shows sample traces of outward currents (voltage-clamp traces) declining in amplitude when aCC was held at -60 mV, close to the chloride reversal potential. The *Drosophila* GABA receptor, *Rdl*, is blocked by 10 µM Picrotoxin (PTX) (Lee et al., 2003) or 1 mM SR95531 (Gabazine) (Hosie and Sattelle, 1996). We observed that the A23a→aCC connection is fully blocked after bath application of either 10 µM PTX (Figure 4H), or 1 mM Gabazine (Figure 4I).

Thus, we confirm that A23a and aCC are monosynaptically connected via a GABAergic synapse. Among the driver lines targeting A23a tested, *SS04495-Gal4: R41G07-AD; R78F07-DBD* (Kohsaka et al., 2019) is the most specific and reliable.

### A31k is monosynaptically connected to aCC

The connectome identifies A31k as a GABAergic interneuron, synaptically connected to aCC, which delivers proprioceptive feedback to motoneurons (Schneider-Mizell et al., 2016; Clark et al., 2018; Kohsaka et al., 2019). We tested three different drivers to verify A31k→aCC connectivity: *R87H09-Gal4* (Zarin et al., 2019); *R20A03-AD; R87H09-DBD* split Gal4 (Zarin, unpublished); and *R20A03-AD; R93B07-DBD* split Gal4 (also termed *SS04399-Gal4* (Kohsaka et al., 2019)).

Evoked AP firing in aCC firing was not influenced following optogenetic activation of A31k using either *R87H09-Gal4* (6.73 ± 0.81 *vs*. 6.68 ± 0.90 APs, pre LED *vs*. LED, respectively, *n* = 7, *P* > 0.99, repeated measure one-way ANOVA followed by Bonferroni’s post hoc test; post LED = 6.46 ± 0.90 APs, data not shown), or *R20A03-AD; R87H09-DBD* split Gal4 (3.20 ± 0.56 *vs*. 3.53 ± 0.43 APs, pre LED *vs*. LED, respectively, *n* = 5, *P* > 0.99, repeated measure one-way ANOVA followed by Bonferroni’s post hoc test; post LED = 3.73 ± 0.50 APs, Figure 5A). Voltage and current-clamp recordings similarly showed no input to aCC, even in absence of TTX (data not shown). Since the connectome was generated from a first instar larval CNS, we repeated our electrophysiological investigation of *R20A03-AD; R87H09-DBD* split Gal4 at the L1 stage. Again, no change in AP firing recorded from L1 aCC was observed during optogenetic stimulation (from 5.99 ± 0.97 to 5.98 ± 1.14, *n* = 5, *P* > 0.99, repeated measure one-way ANOVA followed by Bonferroni’s post hoc test; post LED = 6.41 ± 0.83 APs, Figure 5B). Voltage-clamp recordings performed at -60mV and -40 mV confirmed the absence of any synaptic inputs to aCC following optogenetic activation (data not shown). To further confirm that this line is unreliable, we repeated our stimulation protocol in L3, this time expressing Chrimson, as previously done for one of the A23a split Gal4 (see above). Again, we obtained strong cholinergic, instead of GABAergic inputs, to aCC, most likely due to expression and activation of Chrimson in other interneurons (Figure S1B).

**Figure 5.**
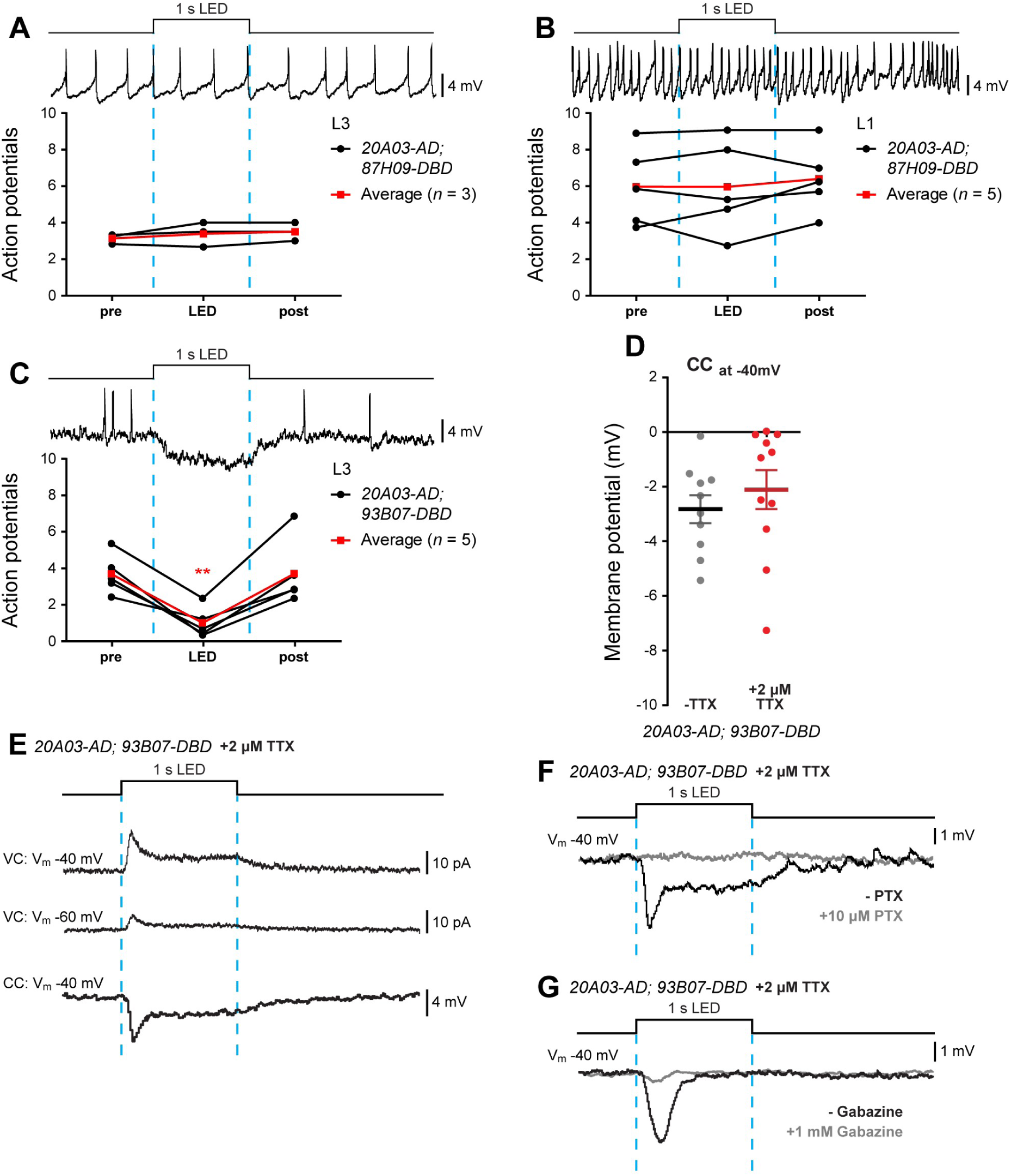
A31k and aCC are monosynaptically connected. (A-B) Optogenetic activation of *R20A03-AD; R87H09-DBD* split Gal4, did not produce detectable changes in aCC firing (evoked by current injection) recorded at both L3 (A: F_(2, 6)_ = 1.733, *P* = 0.3134, repeated measures one-way ANOVA, *n* = 3, black lines), and L1 (B: F_(2, 12)_ = 0.5404, *P* = 0.5928, repeated measures one-way ANOVA, *n* = 5, black lines). Average values are shown in red. (C) Optogenetic activation of *R20A03-AD; R93B07-DBD* split Gal4 significantly reduced aCC firing (F_(2, 12)_ = 20.22, *P* = 0.0011, repeated measures one-way ANOVA, *n* = 5, black lines). Average values are shown in red. (D) Quantification of the synaptic drive to aCC (held at -40 mV) following optogenetic activation of *R20A03-AD; R93B07-DBD* split Gal4, in absence or presence of 2 µM TTX. (E) Sample traces showing the optogenetic activation of *R20A03-AD; R93B07-DBD* split Gal4. aCC neurons were recorded both in voltage (both at -60 and -40 mV) and in current clamp (at -40 mV) in presence of TTX. (F-G) Sample traces confirming the GABAergic connection between A31k and aCC. Cells were recorded, as previously described, before (black trace) and after (gray trace) the bath application of 10 µM Picrotoxin (F), or 1 mM Gabazine (G). In both cases, aCC inputs were abolished.

Optogenetic activation of the split-GAL4 driver line, *SS04399-Gal4: R20A03-AD; R93B07-DBD*, exhibited a clear inhibition of evoked AP firing in aCC (from 3.66 ± 0.49 to 0.99 ± 0.37 APs, *n* = 5, *P* = 0.0088, repeated measures one-way ANOVA followed by Bonferroni’s post hoc test; post LED = 3.68 ± 0.81 APs, Figure 5C). Current-clamp recordings confirmed the presence of a hyperpolarising drive to aCC with no significant reduction after application of TTX (−2.82 ± 0.51 mV, *n* = 10, *vs*. - 2.11 ± 0.72 mV at -40mV, *n* = 11, -TTX *vs*. +TTX, *P* = 0.4347, *t*-test, Figure 5D). Similar to A23a, optogenetic activation of A31k produces an outward current in aCC clamped at -40mV that is reduced at -60 mV (Figure 5E). Again, bath application of 10 µM PTX (Figure 5F), or 1 mM Gabazine (Figure 5G), blocked this inhibitory input to aCC consistent with it being carried by Cl^-^ ions.

These results indicate that A31k makes a GABAergic monosynaptic connection with aCC and that it is selectively targeted by the split Gal4 line *SS04399-Gal4*: *R20A03-AD; R93B07-DBD* (Kohsaka et al., 2019).

## Discussion

In this study we use electrophysiology to validate connectivity of four identified premotor interneurons. Two are cholinergic and form part of the excitatory input to motoneurons, and two are GABAergic and inhibitory (Baines et al., 1999; Rohrbough and Broadie, 2002; Kohsaka et al., 2014; Itakura et al., 2015; Fushiki et al., 2016; Zwart et al., 2016; Kohsaka et al., 2019; Zarin et al., 2019). Collectively, these neurons form part of a central pattern generator that regulates locomotion in *Drosophila* larvae. Indeed, previous recordings from motoneurons indicate that each may receive multiple inputs which has since been validated by connectome data (Baines et al., 1999; Baines et al., 2001; Rohrbough and Broadie, 2002; Zarin et al., 2019).

Our use of NaChBac, which is resistant to TTX, not only provides the required evidence to show monosynaptic connectivity with aCC for these four interneurons (Zhang and Gaudry, 2018), but also reveals useful information about the specificity and reliability of the Gal4 lines adopted in this study (summarised in Table 1). Our results clearly show that this is a powerful method to assess monosynaptic connectivity and likely suitable to be used for verifying other inter-connected cell pairs predicted by the CATMAID dataset in *Drosophila*.

**Table 1.** Summary of results obtained.

| Driver Line | Other names | Interneuron | Connected to aCC | Reference |
| --- | --- | --- | --- | --- |
| R36G02 | A27h-GAL4 | A27h | ✓ | (Fushiki et al., 2016) |
| R15B07 | CLI2-GAL4 | A18a | ✓ | (Zarin , unpublished) |
| R47E12 | CLIs-GAL4 | CLIs | ✓ | (Hasegawa et al., 2016) |
| R47E12; cha3.3-Gal80 | CLI-GAL4 | A18a/A18b3 | ✓ | (Hasegawa et al., 2016) |
| tsh-Gal80; R47E12, cha3.3-Gal80 | CLI1-GAL4 | A18b3 | x | (Hasegawa et al., 2016) |
| SS04495 | R41G07-AD; R78F07-DBD | A23a | ✓ | (Kohsaka et al., 2019) |
| R78F07 |  | A23a | ✓ | (Zarin et al., 2019) |
| No label | R78F07-AD; R49C08-DBD | A23a | x | Zarin, unpublished |
| SS04399 | R20A03-AD; R93B07-DBD | A31k | ✓ | (Kohsaka et al., 2019) |
| R87H09 |  | A31K | x | (Zarin et al., 2019) |
| No label | R20A03-AD; R87H09-DBD | A31K | x | (Zarin, unpublished) |
| Monosynaptically connected to aCC |  |  |  |  |
| No connectivity / lack of specificity |  |  |  |  |

There is a caveat, however, to using this approach due to the significant inactivation of NaChBac at membrane potentials more positive than -40mV (Ren et al., 2001). If a cell expressing NaChBac sat at, or more positive to, this potential then the channel would be almost fully inactivated. Under these conditions, the addition of TTX would block activity completely. The consequent lack of response from the postsynaptic cell could be misinterpreted to mean that two cells are not monosynaptically-connected. Thus, when using this approach, persistence of synaptic drive in the presence of TTX is strong evidence for monosynaptic connectivity, whilst its absence is not definitive. Our data show clearly, in the presence of TTX, that A27h, A18a, A23a and A31k all monosynaptically connect to the aCC motoneuron. Equally, because of the radically different kinetics of NaChBac, compared to the endogenous Na^+^ current mediated by Paralytic, there is little to be gained from analysing the biophysical properties of synaptic drive when this foreign channel is expressed (Baines and Bate, 1998; Baines et al., 2001; Ren et al., 2001). An additional issue with the ectopic expression of an ion channel in a neuron, particularly one of bacterial origin, is that its presence may be sufficient to alter development and/or physiology (Zhang and Gaudry, 2018). Indeed, we show here that expression of NaChBac, either in all cholinergic or GABAergic neurons, is lethal. This technique is also only relevant to synaptic couplings that involve ionotropic receptors, activation of which cause significant change to the postsynaptic membrane potential (and/or Ca^2+^ influx) and neurons that spike (i.e. excludes graded neurons). By contrast, activation of metabotropic receptors, that alter second messenger signalling, are likely to be missed unless ionic movements across the neuronal membrane form part of the activated downstream signalling pathway.

Electrophysiology is a difficult skill for many to master and requires a significant, and costly, amount of equipment. With this in mind, other studies have exploited activity sensors (GCaMP) to replace the requirement for electrophysiology (Sales et al., 2019). Expression of a suitable activity reporter in a presumed postsynaptic cell, together with expression of a cell activator (e.g. optogenetics) overcomes the requirement for the presynaptic cell to support AP firing. Thus, persistence of postsynaptic activity in the presence of TTX is sufficient to infer monosynaptic connectivity. The use of Ca^2+-^ sensors, whilst relative easy to image, can result in a large degree of variability in response, and also normally requires an activation-wavelength that differs to the optogenetic tool being used.

In summary, we validate monosynaptic connectivity for four interneurons with the aCC motoneuron. We find considerable variability between different GAL4 driver lines that purportedly drive in the same premotor neurons. Clearly the question to now address is why? Possibilities include differing expression patterns, differing strengths of expression and/or varying timing of expression. Although for the latter, it should be noted that most of our analysis was carried out at L3 (and also at L1 for selected lines). We are currently imaging these lines to determine where and when they express and will update this preprint as that information becomes available.

## Acknowledgements

The authors would like to thank Chris Doe for generously providing fly stocks and for commenting on this manuscript. This work was supported by funding from the Biotechnology and Biological Sciences Research Council to R.A.B. (BB/N/014561/1) and to M.L. (BB/R016666/1), MEXT/JSPS KAKENHI grants to H.K. (20K06908) and to A.N. (20H05048 and 19H04742) and by a Wellcome Trust Joint Investigator Award to R.A.B and M.L. (217099/Z/19/Z). Work on this project benefited from the Manchester Fly Facility, established through funds from the University of Manchester and the Wellcome Trust (087742/Z/08/Z).

---

**Figure S1.**
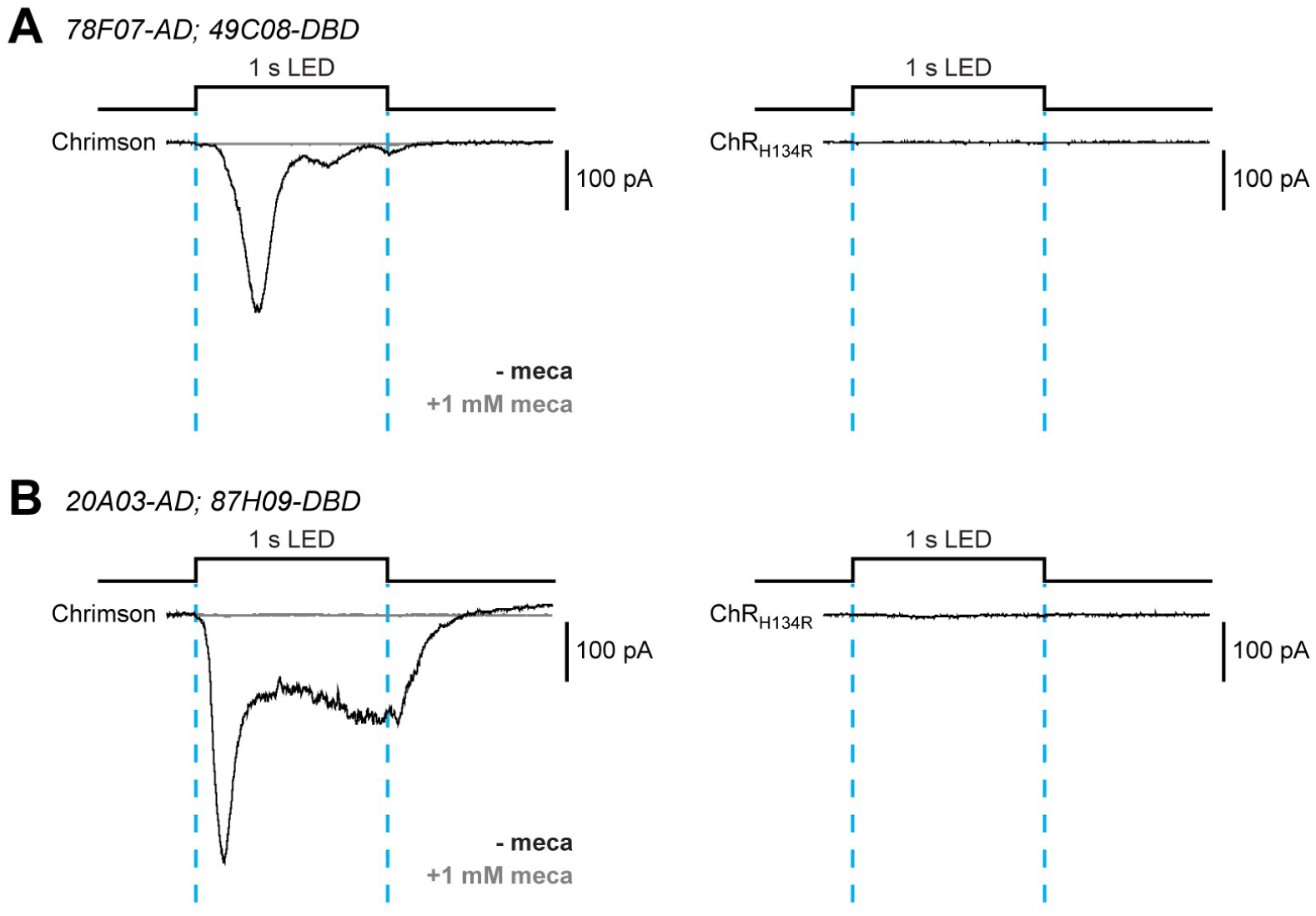
Optogenetic stimulation of Chrimson could generate misleading results. (A) Synaptic drive to aCC obtained by optogenetic activation of *R78F07-AD; R49C08-DBD* split Gal4 driving expression of Chrimson, an extremely powerful ChR variant (Klapoetke et al., 2014). Activation of Chrimson produced an unexpected excitatory inward current in aCC (trace is an average of *n* = 7 cells recorded) which was completely blocked by 1 mM mecamylamine (meca), a commonly used cholinergic blocker. By comparison, expression of H134R-ChR, a variant weaker than Chrimson, showed no detectable inputs (*n* = 10 cells). (B) Synaptic drive to aCC obtained by optogenetic activation of *R20A03-AD; 87H09-DBD* split Gal4 driving Chrimson. Similar to panel A, we observed a mecamylamine-sensitive cholinergic inward current into aCC (*n* = 5 cells) by stimulating Chrimson, while H134R-ChR generated no detectable inputs (*n* = 10 cells).

